# Trans-Acting Genotypes Drive mRNA Expression Affecting Metabolic And Thermal Tolerance Traits

**DOI:** 10.1101/2023.01.15.524165

**Authors:** Melissa K. Drown, Marjorie F. Oleksiak, Douglas L. Crawford

## Abstract

Evolutionary processes driving physiological trait variation depend on the underlying genomic mechanisms. Evolution of these mechanisms depends on whether traits are genetically complex (involving many genes) and how gene expression that impact the traits is converted to phenotype. Yet, genomic mechanisms that impact physiological traits are diverse and context dependent (*e*.*g*., vary by environment or among tissues), making them difficult to discern. Here we examine the relationships between genotype, mRNA expression, and physiological traits to discern the genetic complexity and whether the gene expression effecting the physiological traits is primarily cis or trans-acting. We use low-coverage whole genome sequencing and tissue specific mRNA expression among individuals to identify polymorphisms directly associated with physiological traits and expressed quantitative trait loci (eQTL) driving variation in six temperature specific physiological traits (standard metabolic rate, thermal tolerance, and four substrate specific cardiac metabolic rates). Not surprisingly, there were few, only five, SNPs directly associated with physiological traits. Yet, by focusing on a select set of mRNAs belonging to co-expression modules that explain up to 82% of temperature specific (12°C or 28°C) metabolism and thermal tolerance, we identified hundreds of significant eQTL for mRNA whose expression affects physiological traits. Surprisingly, most eQTL (97.4% for heart and 96.7% for brain) of eQTL were trans-acting. This could be due to higher effect size or greater importance of trans *versus* cis acting eQTLs for mRNAs that are central to co-expression modules. That is, we may have enhanced the identification of trans-acting factors by looking for SNPs associated with mRNAs in co-expression modules that are known to be correlated with the expression of 10s or 100s of other genes, and thus have identified eQTLs with widespread effects on broad gene expression patterns. Overall, these data indicate that the genomic mechanism driving physiological variation across environments is driven by trans-acting tissue specific mRNA expression.

**Author Summary:** In the salt marsh killifish *Fundulus heteroclitus* there is amazingly large variation in physiological traits assumed to be under stabilizing selection, which should reduce their variation. To discern the heritability of this physiological variation we took an innovative approach to define the DNA variation that drives mRNA expression linked to physiological variation. This indirect approach revealed many DNA sequence variants associated with physiological variation *via* their effect on mRNA expression. Surprisingly, these changes were not in the mRNAs themselves, but in unlinked distant genes which regulate mRNA expression. That is, the vast majority (>95%) were trans-acting. This is surprising because trans-acting effects are found less often than DNA variants within or close to mRNA expression genes. Our results are likely related to the select subset of mRNAs across environments that are linked to physiological variation.

## Introduction

For many complex physiological traits multiple genetic loci contribute small effects to produce a continuous phenotypic distribution (Gibson 2010; Bernatchez 2016; Boyle et al. 2017). Some traits have been well studied and the polygenic basis established, including human height (Yang et al. 2011; Turchin et al. 2012; Berg and Coop 2014) and egg production in *Drosophila* and chickens (Szydlowski and Szwaczkowski 2001; Jha et al. 2015). Nevertheless, even when complex physiological traits have substantial heritable physiological variation, their genetic basis often is not as well understood (Maher 2008; Zuk et al. 2012; Simons et al. 2018; López-Cortegano and Caballero 2019). For example, metabolism varies by two to three-fold within populations and by orders of magnitude among species (Burton et al. 2011; Pettersen et al. 2018). Some of this variation can be explained by allometric scaling (relationship to body mass) and environment (Burton et al. 2011; Jayasundara et al. 2015; Schulte 2015; Auer et al.M 2016; Baris et al. 2016; Pettersen et al. 2018); however, after accounting for these and other covariates, the unexplained heritable variation remains high (Bacigalupe et al. 2004; Ronning et al. 2005; Ronning et al. 2007; Nilsson et al. 2009; Wone et al. 2009). Unexpected and diverse molecular and genetic underpinnings have been identified in other complex traits including thermal tolerance (Healy et al. 2018; Drown et al. 2022), brain size (Zwarts et al. 2015; Hoglund et al. 2020), cardiac cellular ATP production (Baris et al. 2017), and flowering time (Andres and Coupland 2012; Frachon et al. 2017; Grabowski et al. 2017). Thus, the relationships between phenotype and genotype for complex physiological traits are multifaceted and likely to be affected by unfamiliar or unexpected genes (Drown et al. 2022). Moreover, physiological traits are context dependent and often vary in different environments or tissues (Jayasundara and Somero 2013; Baris et al. 2016; Chung et al. 2017; Kellermann et al. 2019; Drown et al. 2021). These attributes make it difficult to predict or identify the genetic variation driving physiological variation. One approach to simplify this multifaceted complexity is to identify the genomic mechanism affecting mRNA expression that drives phenotypic variation.

mRNA expression variation is often biologically important in that complex or multivariate mRNA expression can explain variation in a diverse suite of traits including thermal tolerance, disease response, and metabolism (Zhang et al. 2019; Huang et al. 2020; Campbell-Staton et al. 2021; Traylor-Knowles et al. 2021; Drown et al. 2022). Some of this expression is physiologically induced; yet mRNA expression also is heritable (Gibson and Weir 2005) and has large variation among common gardened individuals (Oleksiak et al. 2002). Thus, it is possible to identify heritable genetic loci associated with mRNA expression variation. These associations between genetic loci and mRNA expression are identified as expression quantitative trait loci (eQTL), where eQTLs mediate the expression of one or many genes. Furthermore, when eQTL controlled mRNA expression impacts physiological traits, eQTLs may be evolutionary targets for adaptation (Whitehead and Crawford 2006).

Here, we apply association studies to identify genetic loci directly (genome wide association, GWAS) and indirectly (eQTL) driving physiological variation. We use three common gardened wild populations to capture natural variation in six complex physiological traits: whole animal metabolism (standard metabolic rate, SMR), critical thermal maximum (CT_max_), and four substrate specific cardiac metabolic rates (MR_cardiac_) (Drown et al. 2021). Physiological traits and heart and brain mRNA expression were quantified under two ecologically relevant acclimation temperatures (12°C and 28°C) in the same individuals. We found little evidence of population divergence in physiological traits (Drown et al. 2021) or differentially expressed mRNAs (Drown et al. 2022). Therefore, we treated all individuals as belonging to a single population and, using a tissue specific (heart and brain) weighted gene co-expression network analysis (WGCNA, (Langfelder and Horvath 2008; Healy et al. 2018)), found mRNA expression that explained a large proportion of physiological trait variation (Drown et al. 2022). Data from both studies (Drown et al. 2021; Drown et al. 2022) suggest that variation in these physiological traits is driven by both physiological plasticity and heritable genetic variation among individuals. Whole genome sequencing was used to identify SNPs among the same individuals that were used to quantify physiological and mRNA expression variation allowing us to integrate whole animal (SMR, CT_max_), tissue (MR_cardiac_), and molecular (mRNA expression) level phenotypes (Fig. 1). Specifically, we address four key questions: 1) are SNPs associated with physiological trait variation (direct drivers), 2) are mRNA expression patterns under genetic control (eQTLs), 3) does genetic control of mRNA expression impact physiological trait variation (indirect drivers), and 4) are direct and indirect control mechanisms unique or shared among physiological traits? Few studies have integrated data across levels of biological organization in wild populations to address these questions (Morgante et al. 2020), limiting our understanding of genotypic and molecular variation that impacts complex physiological traits. Using this integrative approach, we find that much of the natural variation in complex physiological traits are affected by trans-acting eQTLs.

**Figure 1:**
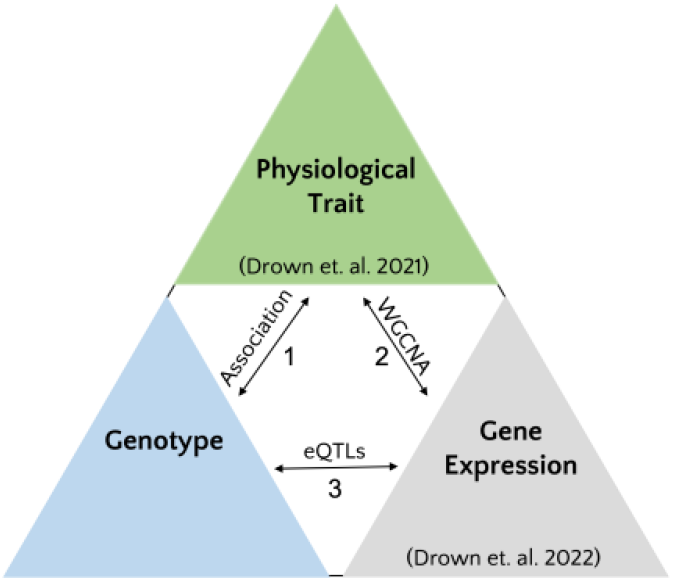
Integrating molecular and genotypic data to understand variation in physiological traits. Physiological trait variation can be driven directly or indirectly (through gene expression) by genotype. To understand the molecular and genetic basis of physiological trait variation, comprehensive datasets can be used to investigate 1) direct associations between genotype and physiological traits, 2) direct correlations between gene expression and physiological traits, for example using weighted gene co-expression analysis (WGCNA), and 3) indirect effects of genotype on physiological traits, which may occur when expression quantitative trait loci (eQTL) impact expression of genes underlying physiological traits.

## Results

### Whole Genome Sequencing results

A total of 172 adult individuals were collected in Fall 2018 (F18) from three geographically close (<15 km) populations, and these were individually barcoded and sequenced to an average depth of 4.1x using low-coverage whole genome sequencing (lcWGS). After data processing and filtering (see methods) 1,406,282 high probability variant sites remained (single nucleotide polymorphisms, SNPs).

### Population structure

To determine the genetic structure among populations, we conducted an admixture analysis. Using NGSadmix we tested seven K values where K is the number of ancestral populations. We found K=4 to be the best fit based on log-likelihood probability with no clear structure among populations (Fig. S1). These analyses, as well as physiological (Drown et al. 2021) and mRNA analyses (Drown et al. 2022) indicate that there is little demographic structure that could affect mRNA expression, physiological traits, or SNPs.

### Linkage disequilibrium

To correct for autocorrelation among SNPs contributing to physiological trait and mRNA expression variation, linkage among SNPs was examined using ngsLD (v1.1.0). Similar to prior studies in this species (Dayan et al. 2019; Ehrlich et al. 2020), linkage among sites decayed within 500 bp with an average R^2^ below 0.2 within 150 bp and below 0.1 within 300 bp (Fig. S2). Thus, only SNPs associated with the same trait that were >500 bp apart were maintained as independent SNPs not in linkage, and those within 500 bp were pruned to keep the most significant SNP. Few SNPs that were significant in the association tests were in linkage and pruning the SNPs for LD did not substantially change the results.

### Association studies

We interrogated potential associations between 1,406,282 SNPs and eight phenotypes: six physiological traits: SMR (standard metabolic rate), CT_max_ (critical thermal maximum), four substrate specific MR_cardiac_ (glucose, fatty acids, LKA, and endogenous), and two measures of mRNA expression: single mRNA and multivariate mRNA expression. Multivariate mRNA expression used weighted gene co-expression network analysis (WGCNA, (Langfelder and Horvath 2008)) to identify co-expression mRNAs and group them into modules (MEs). Single mRNA expression was limited to the top 10 mRNAs from each WGCNA co-expression module. There were 80 MEs: 39 heart modules and 41 brain modules (Table S1), 390 single heart mRNAs, and 410 single brain mRNAs (Drown et al. 2022).

All association results are reported from ANGSD score test (-doAsso 2) and after multiple test correction of p-values (Benjamini and Hochberg 1995). Physiological traits and mRNA expression were measured at two acclimation temperatures (12°C and 28°C). Acclimation temperature was included as a variable in the association tests and yielded similar results when compared to within temperature associations. Sample size varied among association tests based on phenotypic and genotypic data availability and are reported in Table S2.

#### 1. Direct genotypic association to whole animal and tissue level metabolic and thermal olerance traits

A total of five independent SNPs were significantly associated with three traits: GLU MR_cardiac_, LKA MR_cardiac_, and END MR_cardiac_ (Table 1, Fig. 2). Of these five significant associations, two were associated with GLU MR_cardiac_, one was associated with LKA MR_cardiac_, and the remaining two were associated with END MR_cardiac_ (Table 1, Fig. 2). Only one of these SNPs (erfl1, directly associated with END MR_cardiac_) is also a significant eQTL. We did not find significantly associated SNPs for SMR or CT_max_, however, these traits were previously associated with at least one mRNA co-expression module (Drown et al. 2022), described below. The low number of SNPs directly associated with physiological traits is most likely due to the small sample size: average sample size was 29.60 ± 6.23 (mean ± one standard deviation) individuals per association test (Table S2).

**Table 1:** Single nucleotide polymorphisms significantly associated with physiological traits.

| Chromosome | SNP location | Trait | FDR P-value | SNP Annotation | Sample Size |
| --- | --- | --- | --- | --- | --- |
| NC_046380.1 | 38698336 | GLU MR<sub>cardiac</sub> | 2.970000e-08 | foxp1b | 39 |
| NC_046371.1 | 29986564 | GLU MR<sub>cardiac</sub> | 3.150000e-17 | NA | 23 |
| NC_046381.1 | 16773684 | LKA MR<sub>cardiac</sub> | 4.570000e-21 | kcnb2 | 31 |
| NC_046363.1 | 31037486 | END MR<sub>cardiac</sub> | 3.184801e-02 | erfl1 | 25 |
| NC_046371.1 | 19082580 | END MR<sub>cardiac</sub> | 4.859420e-02 | NA | 30 |

**Figure 2:**
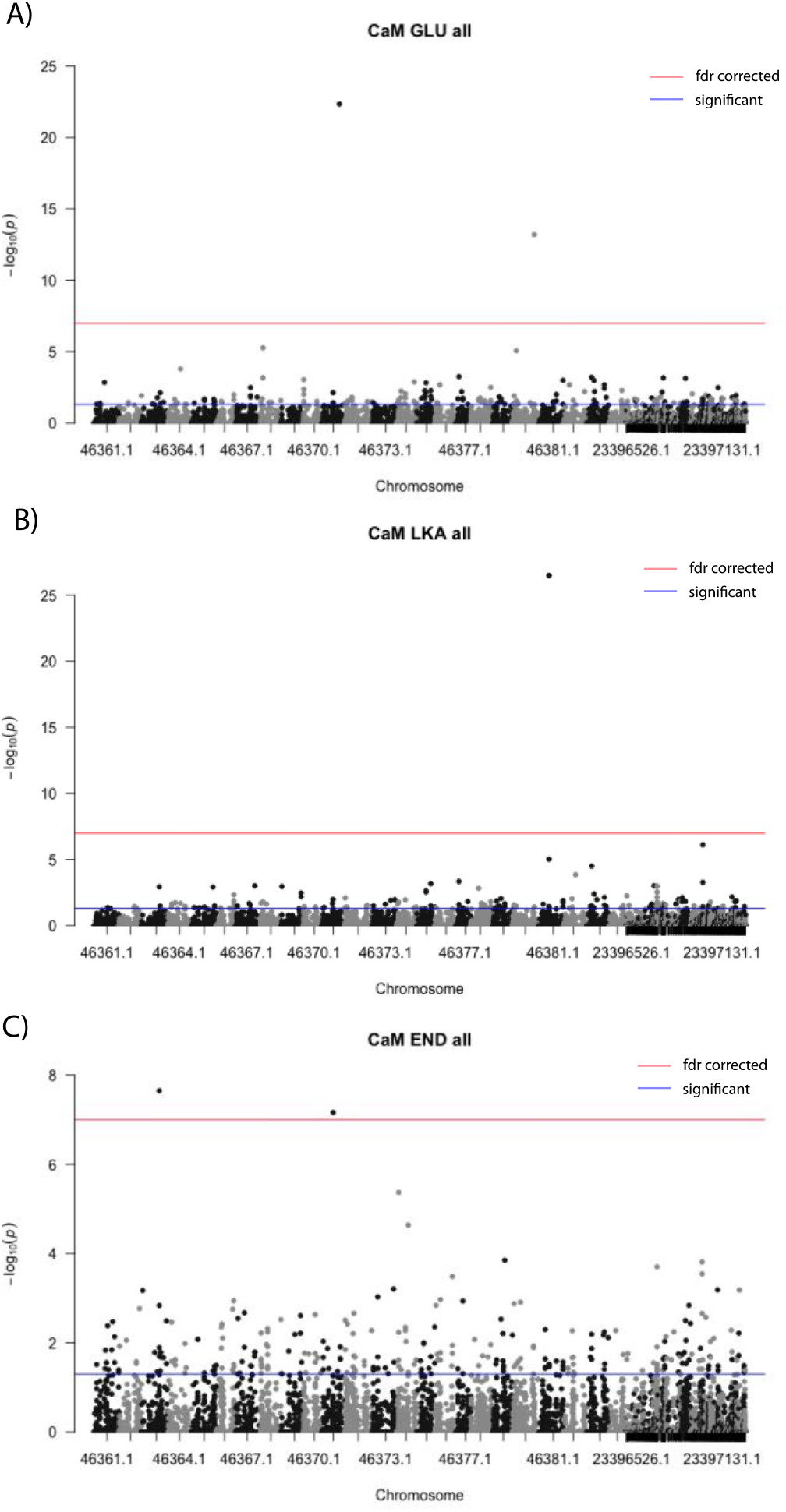
Direct Associations: Manhattan plot for SNPs associated with Cardiac Metabolic Rate. Variation in three substrate specific cardiac metabolic rates (substrates: GLU = glucose, LKA = lactate, ketones, and ethanol, and END = endogenous [no substrate added]) were associated with a total of five single nucleotide polymorphisms (SNPs). **A)** Two SNPs for CaM GLU, **B)** one SNP for CaM LKA, and **C)** two SNPs for CaM END. The five SNPs significant after Benjamini-Hochberg p-value correction are shown above the red line

#### 2. Multivariate and single mRNA expression

We previously identified tissue specific mRNA co-expression modules for heart and brain that are associated with variation in the six physiological traits (Drown et al. 2022). Co-expression modules included 39 significant heart modules with 90-554 mRNAs per module and 41 significant brain modules with 142-393 mRNAs per module (Drown et al. 2022). Each module was assigned a module eigengene (ME, the first principal component of multivariate mRNA expression), and each mRNA in the module was assigned a module membership defined as the correlation coefficient between that mRNA and the ME. From each module we choose the top 10 single mRNAs (based on module membership) and used these single mRNA expression values in a series of association tests to identify eQTL. Additionally, we used the ME for each module as a phenotypic value in a separate set of association studies to find SNPs associated with multivariate mRNA expression (eQTL_ME_). This allowed us to identify SNPs that explain mRNA expression patterns previously correlated to acclimation temperature specific physiological traits (SMR, CT_max_, and MR_cardiac_ measured at 12°C and 28°C) (Drown et al. 2022). Notably, we did not test all possible mRNAs (∼10,000 mRNAs per tissue); instead, we were interested in discrete relationships between SNPs and specific mRNAs and MEs.

### Single mRNA expression associations

For hearts, the 390 single heart mRNAs tested had 79 significant independent genetic associations among 56 unique mRNAs with 52 unique SNPs (Fig. 3). These 79 significant associations with 56 mRNAs and 52 SNPs occur because a SNP tends to be associated with expression of multiple mRNAs (average 1.58 ± 0.95, max=5) and an mRNA tends to have more than one eQTL (average 1.48 ± 0.97, max=4). For brains, the 410 single brain mRNAs tested had 245 significant independent associations among 152 unique mRNAs with 146 unique SNPs (Fig. 3). Again, many SNPs were correlated to more than one mRNA (average 1.68 ± 1.08, max=8), and most mRNAs had more than one significant eQTL (average 2.94 ± 2.49, max=14). Despite testing a similar number of heart and brain mRNAs, brain had 3.1x more total significant eQTL associations than heart (245 brain vs. 79 heart). There was also a greater proportion of mRNAs with at least one significant eQTL in brain (152/410 [37.10%] compared to heart mRNAs (56/390 [14.36%]) (Fig. 3). This difference is not explained by a difference in sample size (i.e., power), which was similar between tissues.

**Figure 3:**
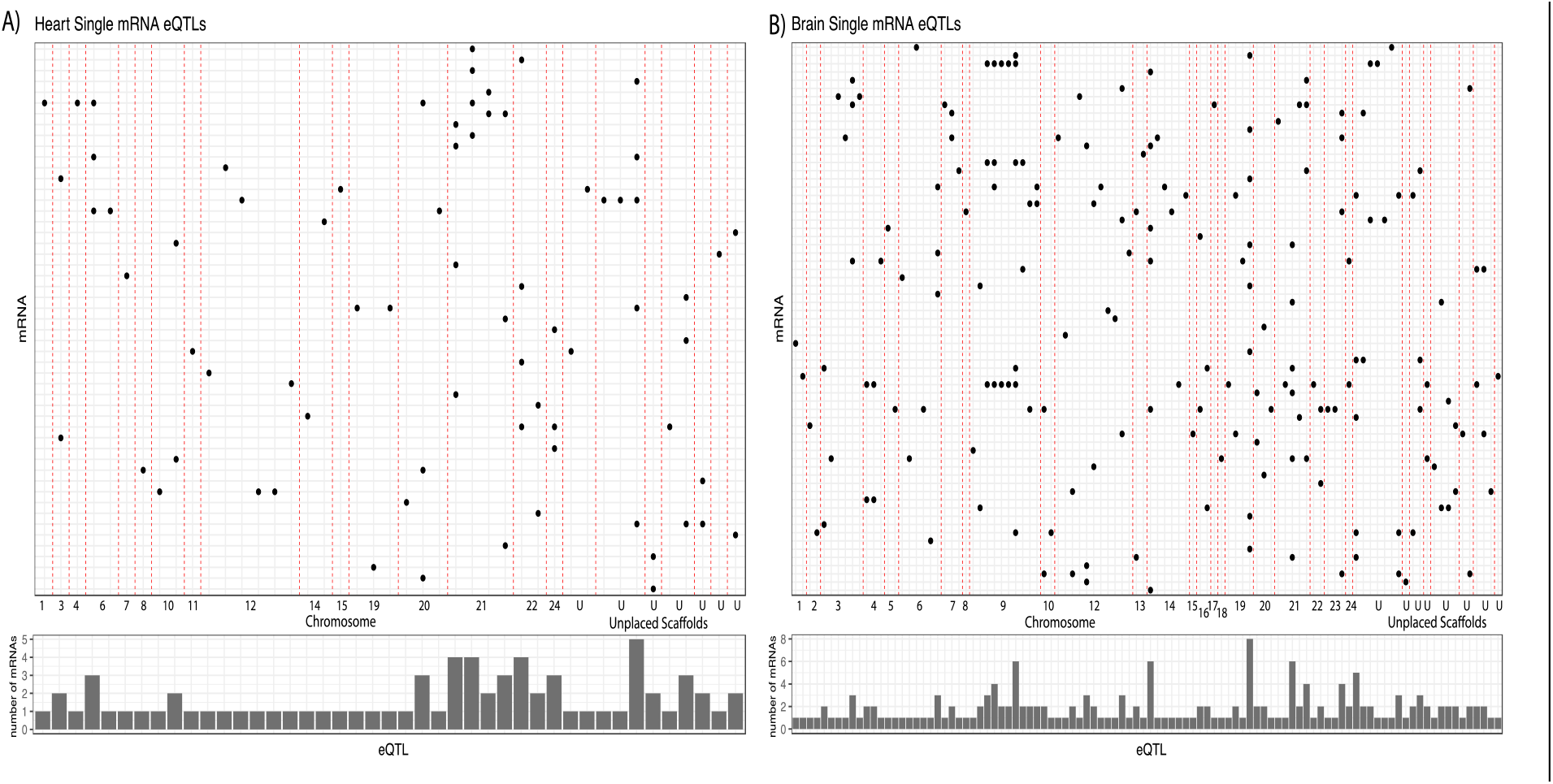
Indirect Associations: Expression quantitative trait loci (eQTL) explain expression of multiple mRNAs in heart and brain. eQTLs for single mRNAs in **A)** heart and **B)** brain tissue. Each SNP that is associated with expression of a mRNA (y-axis) is shown on the x-axis (sorted by SNP position along each chromosome or scaffold, designed by vertical red dashed lines). The bar plots show the number of mRNAs associated with each SNP eQTL (average 1.72 ± 1.01 correlations per SNP for heart [max = 5], average 1.81 ± 1.28 correlations per SNP for brain [max = 8]), sorted by position along each chromosome or scaffold (left to right). All associations are significant with a multiple test corrected p-value < 0.05 (Benjamini and Hochberg).

One explanation for a single SNP being associated with multiple mRNAs is that the mRNA affected by the SNP regulates the expression of many genes. This could occur when an eQTL is found in a transcription factor protein or its regulatory region (e.g., promoter). To determine if an eQTL impacted many mRNAs through a transcription factor, we annotated SNPs and identified those within 5kb of a transcription factor (Table S4). Most of the changes were not in transcription factors. Instead, of the seventeen heart eQTLs (33.7%) associated with more than one mRNA (hereafter identified as “hotspots”), sixteen were in known protein coding regions (within an intron or exon) but only four of these sixteen (25%) were within an annotated transcription factor (erfl1, tada1, atf1, and zbtb3). The one eQTL hotspot not within a known protein coding region was intergenic and not within 5kb of an annotated transcription factor. In brains, 63 eQTLs (43.2%) were identified as hotspots, and of those, 32 were in known protein coding regions (within an intron or exon); and only 2 of the 32 (6.25%) were within an annotated transcription factor (erfl1 and zbtb3) while 1 intergenic SNP was also within 5kb of a zinc finger protein (oocyte zinc finger protein XlCOF6-like).

### Multivariate mRNA expression associations

Among the 39 heart co-expression modules, there were a total of 189 significant associations between 102 eQTL_ME_ and 33 MEs (Fig. 4). These MEs are independent (not correlated to each other) and do not share any mRNAs. As with the single mRNA associations, the relationship between SNPs and MEs was not one to one. A given SNP was correlated with up to six heart MEs with an average of 1.85 ± 1.18 associations per SNP. Of the 102 unique SNPs, 54 were associated with a single heart ME, and the other 48 associated with two or more heart MEs. Among those SNPs correlated with two or more heart MEs, the average number of MEs correlations per SNP was 2.81 ± 1.10.

**Figure 4:**
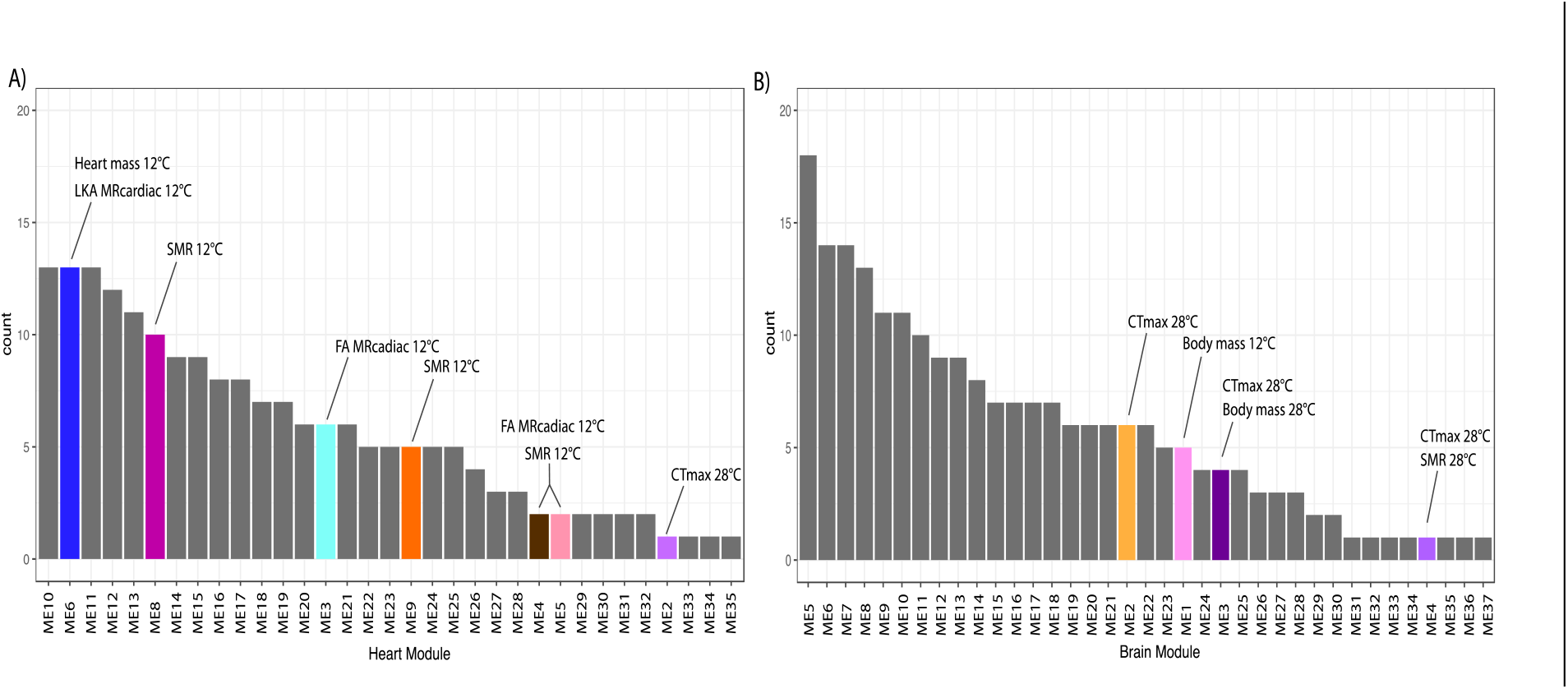
Distribution of multivariate mRNA co-expression associated eQTLs among modules that explain physiological traits. Number of significant SNPs (y-axis, count) associated with each mRNA co-expression module in **A)** heart and **B)** brain tissue. Modules are tissue specific and were identified using a weighted gene co-expression network analysis. Modules previously found to be associated with physiological traits have bright colors and are labeled. Most modules were associated with >1 SNP with an average of 5.72 ± 3.84 SNPs per module for heart (max = 13) and 5.74 ± 4.35 SNPs per module for brain (max = 18).

Among the 41 brain co-expression modules, we found a total of 218 significant associations between 133 eQTL_ME_ and 37 MEs. Single SNPs were correlated with up to six brain MEs with an average of 1.64 ± 1.06 associations per SNP (Fig. 4). Of the 133 unique SNPs, 87 were associated with a single brain ME, and the other 46 associated with two or more brain MEs. Among those SNPs correlated with two or more brain MEs, the average number of ME correlations per SNP was 2.85 ± 1.01. Sample size was similar between heart and brain ME associations.

Similar to the single mRNA eQTL results, several eQTL_ME_ were identified as potential eQTL_ME_ hotspots because they were associated with expression of two or more MEs (Table S5). Out of 48 heart eQTL_ME_ hotspots, 26 were in known protein coding regions (within an intron or exon) with two in annotated transcription factors (jdp2 and atf1). None of the 22 intergenic eQTL_ME_ heart hotspots were within 5kb of an annotated transcription factor. Out of 46 eQTL_ME_ brain hotspots, 21 were in known protein coding regions (within an intron or exon) with two in transcription factors (brf1b, trrap). Additionally, four intergenic eQTL_ME_ were within 5kb of an annotated transcription factor (npas3, XlCGF7.1, and two SNPs in zmym3). One heart and one brain hotspot were also within a gene involved in chromatin remodeling (chd3).

The eQTL studies for single and multivariate mRNAs indicated significant heritable variation underlying tissue specific mRNA expression. Previously, multivariate mRNAs were correlated to physiological traits (Drown et al. 2022). Here, we find that these multivariate mRNA expression patterns are controlled by heritable variation. Thus, many eQTL_ME_ are indirect drivers of physiological trait variation. Specifically, eQTL_ME_ underlying heart mRNA expression are linked to 28°C CT_max_, 12°C FA MR_cardiac_, 12°C SMR, 12°C LKA MR_cardiac_, and 12°C heart mass. Multivariate brain mRNA expression linked to 12°C body mass, 28°C CT_max_, and 28°C SMR is also under indirect genetic control via eQTL_ME_.

### Trans-acting effects on mRNA expression

Although many (61.3% of eQTL and 50% of eQTL_ME_) eQTL and eQTL_ME_ were within genic regions (introns or exons), few (7.5% of eQTL and 6.4% of eQTL_ME_) were found in or near (within 5kb) transcription factors. This is similar to the percentage of transcription factors found globally in eukaryotic genomes and indicates that there is not an enrichment for eQTL or eQTL_ME_ within or near transcription factors (Riechmann 2002). However, the large number of eQTL and eQTL_ME_ (50-61.3%) found within genic regions (introns or exons) exceeds expectations (∼2% of eukaryotic genomes are protein coding). Prior studies have reported an enrichment within and near genic regions for trait associated SNPs (Li et al. 2012; Watanabe et al. 2019) and a high likelihood of trait associated SNPs being eQTLs (Nicolae et al. 2010).

To better understand the genomic context of eQTL and eQTL_ME_ we determined their proximity to the mRNA(s) they effect. Interestingly, we found that >95% of single mRNA eQTLs (97.4% for heart and 96.7% for brain) were trans-acting (defined as SNPs found on a different chromosome or scaffold than that of the mRNA with which they were associated). Specifically, for heart 94.1% (16/17) of eQTL hotspots, and 97.1% (34/35) of eQTL impacting single mRNAs were trans-acting. For brain 95.2% (60/63) of eQTL hotspots, and 96.4% (80/83) of eQTL impacting single mRNAs were trans-acting. While eQTL_ME_ could not be classified as cis or trans-acting because they effect co-expression modules containing many disparately located mRNAs, we determined whether the eQTL_ME_ were found within 5kb of mRNAs that were part of the module. All heart and 98% of brain eQTL_ME_ were in genes not containing mRNAs from that module. Thus, eQTL_ME_ are not simply cis-acting on a high-ranking mRNA within the module but have broad effects on the expression of many module mRNAs.

### Patterns of shared association

For both heart and brain, we looked for eQTL association for 10 mRNAs from each co-expression module. Within an ME, the 10 mRNAs have some degree of correlation with each other causing them to be grouped into the same module. Thus, we examined whether mRNAs from the same module had more shared eQTL than those from different modules. Surprisingly, within 83% of heart and 67% of brain modules the top ten mRNAs all had unique eQTLs. This “uniqueness” is demonstrated in the frequency of shared eQTLs within *versus* among modules: for both hearts and brains, there was fewer shared eQTLs within modules than among modules (T-test, heart p-value=0.027, brain p-value=0.008). This eQTL uniqueness would be expected if they were cis acting, for example, a SNP in the mRNA promoter. Yet, many eQTLs are not near or within the genes encoding the mRNA they effect. This indicates that these eQTL are trans-acting regulatory SNPs in the sense that they are found outside of the genes that code for modular mRNAs. Similarly, for eQTL_ME_, nearly all (98% of brain and 100% of heart) were in genes not containing mRNAs from that module. Instead, they were found in different parts of the genome and in genes unassociated with the physiological trait(s) correlated to the module.

Finally, we looked for overlap among SNP sets associated with a given phenotype (physiological traits, tissue specific single mRNAs, or tissue specific MEs, Fig. 5). First, to address whether the eQTL_ME_ were impacting module expression through action on a single high-ranking mRNA in that module, we assessed whether any SNP was associated with an mRNA and the module containing that same mRNA (overlap in eQTL and eQTL_ME_). There were no shared mRNAs between eQTL and eQTL_ME_, suggesting that the eQTL_ME_ are not acting on a single mRNAs but represent a more complex mechanism of multivariate mRNA expression control. Second, we examined overlap between tissues for SNPs associated with both single mRNAs and modules. There was only one mRNA (atp7a, 0.13% of total) explained by at least one heart and at least one brain eQTL. However, the eQTLs for this mRNA were not the same for heart and brain expression. These data suggest that in the different tissues the variation in atp7a mRNA is affected by different SNPs. There is a limited generality of this finding because we only tested single heart and brain mRNAs that were high-ranking in modules, with only 3% of tested mRNAs shared between heart and brain. Thus, while our results suggest that genetic control of mRNA expression is tissue specific, examining a larger set of mRNAs expressed in several tissues would likely be more informative about the role of conserved eQTL among tissues.

**Figure 5:**
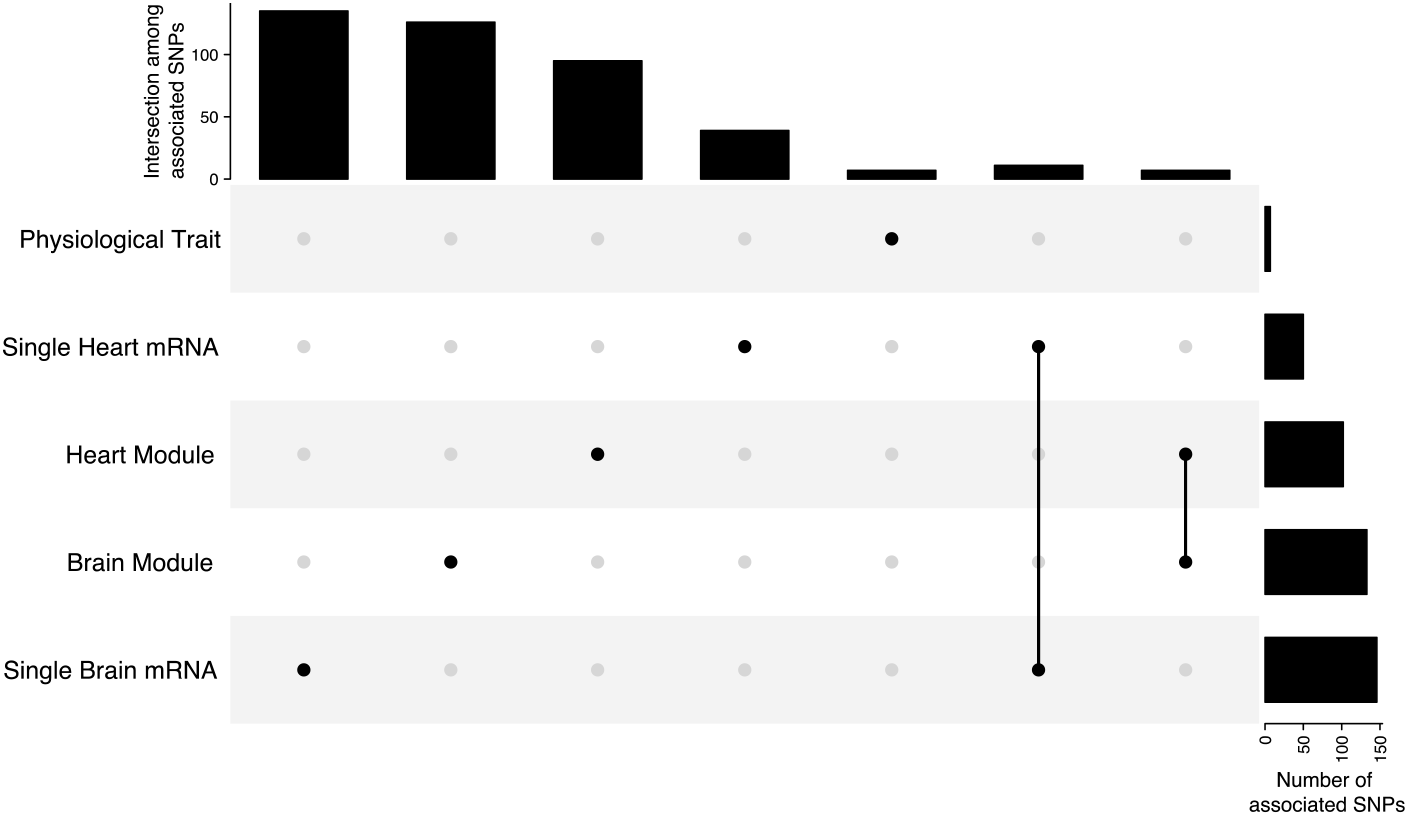
Unique eQTLs explain single mRNA and multivariate mRNA co-expression between heart and brain tissues. Overlap among eQTLs associated with single mRNAs and co-expressed mRNA modules (Heart and Brain Modules) and SNPs associated with physiological traits. In both tissues, SNPs uniquely explain either single or co-expressed mRNAs with few shared among SNP sets (five rightmost vertical bars). Twelve (6.2%) eQTLs are correlated with single mRNA expression in both heart and brain tissues and seven (3.0%) eQTLs are correlated with module expression in both heart and brain tissues. Finally, a single SNP (chr NC046363.1 location 31037486) was correlated to expression of lmnb1 (a protein involved in cellular architecture) in brain and endogenous cardiac metabolism. Horizontal bars (far right) show the number of SNPs associated with each variable (single mRNA expression, module mRNA expression, or physiological traits), vertical bars (top) show number of SNPs unique (five leftmost vertical bars) or shared (five rightmost vertical bars) among datasets.

### Genetic diversity

To compare genetic diversity among SNP sets, heterozygosity (H_e_) was quantified. Among all 1,406,282 high-probability variant sites, the average heterozygosity was 0.231 ± 0.164 (mean ± standard deviation). Heterozygosity for eQTL SNPs for heart and brain were significantly higher than for all SNPs (heart = 0.328 ± 0.074, brain = 0.325 ± 0.068) and was highest for eQTL_ME_ SNPs (heart = 0.368 ± 0.091, brain = 0.361 ± 0.099). Tissue specific eQTL and eQTL_ME_ SNPs did not significantly differ in heterozygosity (Fig. S3).

## Discussion

We examined associations among SNPs and specific mRNAs that were previously identified as biologically important based on their membership in co-expression modules. This reduced our test from ∼1.5 million SNPs across ∼10,000 mRNAs in each tissue to <500 mRNAs in each tissue. Additionally, we use WGCNA (Langfelder and Horvath 2008) to summarize multivariate mRNA expression for co-expressed mRNAs. Using this approach, we summarized ∼10,000 mRNAs per tissue into 39 heart and 41 brain modules that could be used in our association tests. In comparison to testing each of these mRNAs individually this may have reduced the signal to noise ratio in our mRNA expression data and increased our power by reducing the number of tests (Westra and Franke 2014).

The data suggest that there are many more significant indirect eQTL than direct SNP associations for the physiological traits we examined. Most of these indirect eQTL were trans-acting and in known protein coding regions (within introns or exons). Many (33.7% of heart and 43.2% of brain) eQTL were hotspots--associated with more than one mRNA. These results are affected by the power of our analyses. While we used a total of 172 individuals, a minority of individuals (∼25) had genotypic data at any given loci limiting the sample size of the association tests. While other association studies conducted in wild populations use a similar number of individuals (Hecht et al. 2013; Bourret et al. 2014; Scott et al. 2015; Campbell-Staton et al. 2021), we acknowledge that this may limit our findings in two ways. First, small sample sizes could limit findings to only large effect loci. Yet, the important insights are that nearly all significant eQTL are trans-acting and affect multiple mRNAs that are linked to physiological variation. The observation that nearly all significant eQTL are trans-acting suggests that trans-acting eQTL have larger effect size than cis acting eQTL, which allowed us to detect them here. Second, small sample sizes may increase the risk of detecting spurious associations. Yet, the observation that eQTL were trans-acting and affecting multiple mRNA indicates that the eQTL are not spurious. This is, both because it is unlikely that multiple independent associations would spuriously be significant for a given eQTL and because it is unlikely that trans-acting factors (defined as being on one of the 24 separate chromosomes) would represent more than 90% of significant associations if they were spurious. Thus, while all our conclusions are based on the limits of detections, these limits indicate the importance of trans-acting factors affecting physiologically important mRNA expression. Still, we caution against making assumptions about the role of specific SNPs that have been identified here in explaining physiological traits or gene expression in other populations. This is both due to the limited sample size and the lack of direct evidence (e.g., a traditional QTL study) for potential quantitative trait loci and eQTLs. Instead, we focus the discussion on general patterns that speak to the genetic and molecular control of physiological traits.

Our first question asked if there were SNPs directly associated with physiological trait variation. We found only five SNPs directly association with physiological trait and all five were to substrate specific MR_cardiac_. Second, we asked if mRNA expression patterns were under genetic control. We found many significant eQTL associated with single mRNAs or MEs in heart or brain tissue suggesting that mRNA expression patterns are influenced by genotype and likely heritable (Table S2). Third, we leveraged data from our prior study that found significant correlations between mRNA expression modules and physiological traits to ask if genetic control of mRNA expression was likely to impact physiological trait variation. Nine heart modules and four brain modules were previously identified as biologically important (correlated to physiological traits). Of these, seven out of nine heart and all four brain modules were associated with at least one eQTL_ME_. Our results suggest that physiological traits are heritable *via* genetic control of gene expression, although we note that because of the limited sample size, there are likely more trait associated loci than we have detected here. Fourth and finally, we asked if direct and indirect control mechanisms were unique or shared among physiological traits. That is, we investigated if physiological traits had a shared genetic (direct) or molecular (indirect) basis. Only one direct associated SNP was also an eQTL (erfl1), the remaining eQTLs suggest unique genetic and molecular mechanisms underlying the physiological traits we examined.

We treated individuals as belonging to a single population based on the admixture analysis, which revealed no structure among the three sampled populations (Fig S1). This conclusion is additionally supported by three prior studies: first, we found few differences among populations in SMR, CT_max_, or substrate specific MR_cardiac_ when measured under two ecologically relevant temperatures (12°C and 28°C) (Drown et al. 2021); second, mRNA expression also contained few population specific differences and shared mRNA co-expression modules were significantly associated with physiological trait variation among all individuals (Drown et al. 2022); third, Dayan et. al. found no structure among three similar central New Jersey populations of *F. heteroclitus* (same central site, more geographically distant north and south reference populations) (Dayan et al. 2017). Notably, prior studies have found that mRNA expression explains up to ∼80% of variation in MR_cardaic_ (Oleksiak et al. 2005; Drown et al. 2022) as well as SMR and CT_max_ (Drown et al. 2022). Here, we build on this prior knowledge to identify SNPs associated with physiological traits and eQTLs that explain single and multivariate mRNA expression.

### Direct and indirect control of physiological traits

#### 1. Direct genetic control of physiological traits

Three out of six physiological traits (GLU, LKA, and END MR_cardiac_) were associated with five SNPs (Table 1, Fig. 1). All three of these traits are for cardiac metabolism, potentially reflecting less complex genomic architecture than standard whole animal metabolism or CT_max_. Three of the five SNPs were within annotated genes: foxp1b (GLU MR_cardiac_), kcnb2 (LKA MR_cardiac_), and erfl1 (END MR_cardiac_). Foxp1b is forkhead box protein and erfl1 is a repressor factor; both are involved in negative regulation of transcription by RNA polymerase II (Cheng et al. 2007; Liu and Patient 2008). Kcnb2 is a voltage gated potassium channel, with various functions including but not limited to regulation of neurotransmitter release, heart rate, insulin release, and smooth muscle contraction (Hristov et al. 2012; Li et al. 2022). These three genes are not found in any of the heart or brain mRNA co-expression modules. Yet, one gene (erfl1) directly associated with END MR_cardiac_ contains different SNPs that are eQTL effecting mRNA expression in hearts and brains.

Our study did not find any direct association with *F. heteroclitus’* CT_max_, but a prior study found 47 candidate SNPs associated (Healy et al. 2018). Similarly, none of our SNPs were directly associated with standard metabolic rate while prior studies have found multiple candidate quantitative trait loci (QTL) associated with metabolic rate (Jacobson 2006; Palomar et al. 2019). It is likely, given the complex nature of metabolic and thermal tolerance traits, that many small effect loci rather than one or few large effect loci explain these traits (Wentzell et al. 2007; Burton et al. 2011; Csillery et al. 2018; Healy et al. 2018). In fact, multiple studies provide evidence for a polygenic basis of both traits (Healy et al. 2018; Barghi et al. 2019). Our approach, which is best suited to detect large effect associations is underpowered. Thus, although we detected few direct drivers of physiological trait variation, this is unlikely to be representative of the biology. Using a multivariate approach where genotypes at many SNPs could be used to explain these traits may improve our detection of associated SNPs, as has been done in other studies (Bourret et al. 2014; Healy et al. 2018; Barghi et al. 2019). Instead of this multivariate approach, we focus on mRNA expression patterns that may indirectly impact physiological traits. The ability of this approach to identified indirect drivers (eQTLs for mRNAs linked to physiological traits) may be due to high effect sizes of eQTL compared to QTL, especially for multivariate mRNA expression (Westra and Franke 2014; Boyle et al. 2017). That is, where physiological traits are complex and affected by 100s to 1,000s of loci, mRNA expression is likely to be affected by fewer loci; thus, each eQTL will have larger effect because fewer polymorphisms are involved (Boyle et al. 2017).

#### 2. Are mRNA expression patterns under genetic control?

In both heart and brain tissue, we found eQTLs for 14.6% and 20.7% of the tested heart and brain mRNAs respectively, suggesting that heritable genetic variation is important in explaining these mRNA expression patterns (Fig. 2). Interestingly, the majority (97.4% for heart and 96.7% for brain) of eQTL that we detected were found on different chromosomes than the affected mRNA (trans-acting). This could reflect a higher effect size or greater importance of trans *versus* cis acting eQTLs for the selected mRNAs expression. The relative role of cis and trans-acting factors is often examined; however, many studies have found a prominent role of cis elements in comparison to trans elements (Kitano et al. 2019), although see (Hoglund et al. 2020)). Here, we find the opposite with few (<5%) of all eQTL in heart and brain being cis acting (found on the same chromosome). If trans-acting factors have a larger effect size, this may be explained by our increased likelihood of detecting trans *versus* cis factors; however, other studies often struggle with the detection of trans-acting factors, which is often attributed to their lower effect size (Nica and Dermitzakis 2013; Westra and Franke 2014). Thus, our ability to detect trans-acting factors is unique and may be explained by the selection of specific mRNAs that are central to co-expression modules for our tests. That is, we may have enhanced our ability to identify trans-acting factors by looking for SNPs associated with mRNAs in co-expression modules that are known to be correlated with the expression of 10s or 100s of other genes. Thus, by examining a select set of single mRNAs, we may have captured a large number of transcription factors or other regulatory elements likely to have widespread effects on broad gene expression patterns. As eQTL effects are often context dependent, our examination of tissue and temperature specific gene expression also may have contributed to the number of significant trans-acting elements we have detected (Westra and Franke 2014; Boyle et al. 2017). In contrast, other studies may use the whole transcriptome (rather than specific tissues) and rarely examine mRNA expression under multiple environments. The frequency of trans-acting effects is important because it is often undetected and unappreciated. (Price et al. 2011).

While we did not classify SNPs associated with MEs as cis or trans because there is no single genomic location for a module, we did examine whether SNPs associated with modules were found in genes that produced mRNAs that were part of that module. We found no association between SNP identity and module membership. That is, nearly all the eQTLs for MEs were not found in or within 5,000bp of the genes for the mRNAs in our MEs. Additionally, the SNPs associated with single mRNAs were not the same as those associated with ME expression, suggesting that eQTL_ME_ were functionally independent from any single mRNA. This demonstrates that ME expression is not driven by the effect of an eQTL on a single high-ranking mRNA within the module. No studies to our knowledge have used multivariate mRNAs defined by a WGCNA analysis as a phenotypic outcome in a genetic association study.

The approximately 25 individuals for any one SNP limits conclusion on the diversity of mRNA and molecular mechanisms affecting the six physiological traits. Yet, we find many significant eQTLs for both single and multivariate mRNA expression where a vast majority were trans-acting or distant to the variable mRNA loci. These findings suggest that our approach may allow for better detection of trans-acting elements, which may impact expression of dozens or hundreds of co-expressed mRNAs.

Finally, we find that eQTL and eQTL_ME_ have greater heterozygosity when compared to all 1,406,282 variant sites used in this study. This may be explained by the association approach as less variable sites have less variance among individuals that can be used to explain variance in the physiological traits and mRNA expression patterns. It is also possible that heterozygosity differences are biologically relevant. If eQTL and eQTL_ME_ sites are under directional selection, we might expect a loss of genetic diversity in these sites. Yet, we find that heterozygosity is higher in eQTL and eQTL_ME_ when compared to all SNPs. Higher heterozygosity may be due to genetic redundancy, as indicated by the presence of different SNPs underlying eQTL and eQTL_ME_ and the association of correlated physiological traits with different underlying loci. These patterns could be driven by spatial or temporal variation in selection preventing allelic fixation and aiding in the maintenance of biologically important variation at the SNP and mRNA level.

#### 3. Are genetically controlled mRNA expression patterns likely to impact physiological traits?

Heritable genetic variation impacts mRNA expression (Gibson and Weir 2005), a molecular level phenotype that is well established as important in physiological response to the environment (McCairns and Bernatchez 2010; Albert and Kruglyak 2015; Dayan et al. 2015; Zhang et al. 2019; Campbell-Staton et al. 2021). Thus, our finding of genetic links to mRNA is not surprising, even with small sample size. What is unique in our data is the ability to examine relationships among genotype and gene expression (eQTLs) and interpret them in the context of known correlations between gene expression and physiological traits. This may be especially important for improving our ability to detect biologically important genetic variation (Albert and Kruglyak 2015) because gene expression is a relatively simpler trait that may be impacted by fewer, larger effect, and easier to detect loci (Boyle et al. 2017).

Previously, we found that up to 82% of the variation in several temperature specific (12°C or 28°C) metabolic and thermal tolerance traits could be explained by co-expression among hundreds of mRNAs grouped into modules (Drown et al. 2022). However, we could not parse the roles of plasticity *versus* heritable mRNA expression variation. Here, we show that both expression of single mRNAs found within these modules and multivariate module expression is associated with genetic variation. This suggests that a large portion of several physiological traits may be explained by heritable mRNA expression variation. Specifically, heritable variation in heart mRNA expression is linked to 28°C CT_max_, 12°C FA MR_cardiac_, 12°C SMR, 12°C LKA MR_cardiac_, and 12°C heart mass. Heritable variation in brain mRNA expression is linked to 12°C body mass, 28°C CT_max_, and 28°C SMR. Of the modules previously correlated with temperature specific physiological traits, almost all (77.8% of heart and all brain) had at least one significant association with at least one SNP. This provides further evidence that mRNA expression patterns impacting these physiological traits are under genetic control and heritable.

Prior studies have demonstrated that heritable mRNA expression variation can impact diverse physiological traits. For example, behavioral maturation from hive worker to forager between honey bee subspecies is partially attributed (up to 30%) to heritable variation in brain gene expression (Whitfield et al. 2006). Various organisms including sea turtles (Tedeschi et al. 2016), maize (Frova 1993), fruit flies (Sejerkilde et al. 2003), and fish (Fangue et al. 2006; Heredia-Middleton et al. 2008) among others exhibit heritability of heat shock protein expression, allowing them to respond to environmental temperature variation. Here, we found that biologically important single and multivariate mRNA expression related to physiological traits has a genetic basis and is heritable. This is similar to studies that have found overlap among QTL and eQTL sets (Wentzell et al. 2007; Carrasco-Valenzuela et al. 2019), yet the finding that a single eQTL is significantly associated with 100s of co-expressed mRNAs is unique. Further, many of the modules were associated with more than one eQTL, suggesting that there is substantial genetic variation contributing to gene expression patterns related to physiological traits. This heritable variation in gene expression provides the raw material for evolution and may explain the vast inter-individual variation in physiological traits that we have measured (Drown et al. 2021; Drown et al. 2022).

#### 4. Are physiological traits genetically independent?

Here, we have shown that mRNA expression patterns previously correlated with physiological traits are under genetic control by a suite of mostly trans-acting eQTL. One additional potential outcome of this study was to determine if physiological traits had a shared genetic basis. This was an important avenue to explore as our prior studies found correlation among traits (Drown et al. 2021) and a shared molecular basis among correlated traits (Drown et al. 2022). For example, we previously found that 12°C FA MR_cardiac_ was positively associated with 12°C SMR and that these traits were both associated with two MEs (heart ME4 and heart ME5). If these traits also had a shared genetic basis, we expected eQTL_ME_ for heart ME4 and heart ME5 to be shared. Despite the shared association with these two traits, these MEs were associated with two different eQTL_ME_ each. Similar patterns were found in brain among ME2, ME3, and ME4, which were all correlated with 28°C CT_max_ but did not share any eQTL_ME_. In general, eQTL_ME_ were correlated to more than one ME (46.2% of heart and 35% of brain); however, only four SNPs (1.7%) were shared among >1 module that was also associated with a physiological trait (Fig. S4). Additionally, the five SNPs associated directly with physiological traits were not shared among traits and do not overlap with ane eQTL, although only one gene contains multiple SNPs that are either directly associated with a trait or are eQTLs (erfl1).

Overall, the limited overlap among SNPs associated directly or indirectly (eQTL) with physiological traits within a tissue was surprising. This may be due to our limited ability to detect small effect QTL, and we might expect greater overlap between QTL and eQTL if more mRNAs were tested across more individuals. However, within the power of our data, we detected diverse and complex molecular mechanisms contributing to physiological trait variation. Tissue specific expression patterns appear to be under unique genetic control and multivariate mRNA expression is not explained by a single eQTL impacting mRNA expression of a gene highly correlation to a given module.

Thus, while different traits are correlated to the same ME, the nucleotide polymorphism, or genetic control, of mRNA expression is distinct. This suggests that there is substantial genetic variation underlying the physiological traits we have measured, with a diversity of molecular and genetic mechanisms contributing to trait variation. The paradoxical genetic independence of physiologically related traits (here metabolism and thermal tolerance) is not uncommon (see (Herrewege 1980; Baker et al. 2015; Healy et al. 2018)) and emphasizes that these traits may still be evolutionarily distinct, although they are linked at the molecular or physiological level.

## Conclusions

Relationships among genotype, gene expression, and physiological traits explain biologically important natural variation found in wild populations. In particular, substantial and diverse genetic variation impacts these traits through direct and indirect (eQTL and eQTL_ME_) mechanisms. Demonstrated here, much of the mRNA expression variation is associated with a diverse set of trans-acting eQTLs. Surprisingly, these trans-acting eQTLs are unique even for mRNAs that affect multiple traits. Under a simpler genetic architecture, we may expect mRNAs that have a shared association with cardiac and whole animal metabolism to also share the same trans-acting eQTL, but this does not occur. Instead, the mRNA expression changes that affect multiple physiological traits are regulated by different trans-acting SNPs. Finally, the SNPs directly or indirectly associated with physiological traits have greater heterozygosity (genetic variation) compared to all SNPs, and this greater genetic variation likely contributes to *F. heteroclitus’* well characterized resilience and plasticity (reviewed in (Burnett et al. 2007; Crawford et al. 2020)) in the face of novel environments. Together, our data suggest genetic control of biologically effective, mRNA expression (expression that impacts physiological traits), which in turn, may impact fitness.

## Methods

### 1. Sample collection

Fin clips were taken from adult *F. heteroclitus* collected along the central coast of New Jersey, USA near the Oyster Creek Nuclear Generating Station (OCNGS), which produces a thermal effluent that locally heats the water. Three populations were sampled: 1) north reference (N.Ref; 39°52’28.000 N, 74°08’19.000 W), 2) south reference (S.Ref; 39°47’04.000 N, 74°11’07.000 W) and 3) a central site located between the southern and northern reference that is within the OCNGS thermal effluent (TE; 39°48’33.000 N, 74°10’51.000 W). The TE population used here differs by 4°C in habitat temperature from the two references populations (average summer high tide temperature 28°C N.Ref and S.Ref, and 32°C for TE) but is otherwise ecologically similar (Drown et al. 2021) (Dayan et al. 2019). Fin clips were collected in fall 2015 (F15), fall 2018 (F18), spring 2019 (S19), fall 2019 (F19), and fall 2020 (F20) and stored in GuHCl buffer. DNA was extracted using carboxyl coated magnetic beads. The DNA quality was assayed using gel electrophoresis and spectrophotometry to ensure high molecular weight and low contamination.

### 2. Library preparation

The analysis presented here uses a subset of samples that were part of a larger sequencing run. A total of 1121 individuals were sequenced (Table S3). All samples were quantified in triplicate using spectrophotometry and normalized to 100ng for sequencing library preparation. The whole genome sequencing library was prepared using a tagmentation approach. Briefly, DNA was digested with an in-house purified Tn5 transposase (as in (Picelli et al. 2014)) loaded with partial adapter sequences. After tagmentation, the fragmented DNA was amplified using barcoded primers such that each individual sample would contain a unique i7 and a plate level (1 per 96 samples) i5 barcode. This allowed for unique dual indexing of up to 768 individuals. After barcoding, samples were combined into two pools (560 samples each) and each pool amplified and then sequenced on a single lane of Illumina HiSeq 3000. These single sequencing lanes were assessed to determine coverage balance among samples, and the same libraries were sequenced across an additional four lanes each. For all sequencing runs, a greater relative amount of library for F18 samples was added to the pool to achieve higher coverage because whole animal, tissue, and molecular (mRNA expression) level phenotypic data was available for these individuals.

### 3. Raw sequence analysis

We followed best practices for low coverage whole genome sequencing (lc-WGS) data processing as in (Lou et al. 2021). Briefly, adapter sequences and low-quality bases were trimmed using Trimmomatic (v0.39) (Bolger et al. 2014). Flash (v1.2.11)(Magoč and Salzberg 2011) was used to combine overlapping reads and to parse singletons and paired reads. Singletons and paired reads were mapped separately using BWA mem (v0.7.17) and resulting sam files converted to bam files using samtools (v1.3.1) (Danecek et al. 2021). The first and second sequencing run were processed separately until BAM files were produced and found to be of similar quality assessed by comparing total percentage of mapped reads and levels of dually mapped reads before combining for the remaining file processing steps. Picard (v2.26.4) was used to add read group information, which is needed for duplicate marking downstream. After combining all mapped reads for a single individual, BAM files were further filtered for mapping quality using samtools and overlapping reads softclipped using bamutil (v1.0.15). Finally, Picard (v2.26.4) was used to mark duplicate reads.

### 4. Variant calling

Two variant calling pipelines were used. First, Freebayes (v1.0.2) was used to call variants and the resulting VCF file was filtered using VCFtools (v0.1.16). VCFtools filters were to include only biallelic sites, >5% minor allele frequency, <5% missingness per individual, <10% missingness per site. This resulted in 1,406,282 high-probability variant sites. ANGSD (v0.935) (Korneliussen et al. 2014), which is designed for use with lcWGS data, was then used to obtain a genotype likelihood beagle file containing the previously identified high-probability variant sites from Freebayes and VCFtools. This approach is similar to other studies using lcWGS data where variant calling may be sensitive to specific tool use.

#### Phenotypic data

Methods and analyses for phenotypic data are described in previous publications (DeLiberto et al. 2020; Drown et al. 2020; Drown et al. 2021; Drown et al. 2022). Briefly, all phenotypes were measured after common gardening and under two temperature acclimation conditions. Whole animal phenotypes included temperature specific whole animal metabolism (standard metabolic rate, SMR) and critical thermal maximum (CT_max_, a measure of thermal tolerance) measured at and after acclimation to 12 and 28°C. Tissue level phenotypes included four substrate specific cardiac metabolic rates (MR_cardiac_, substrates: glucose [GLU], fatty acids [FA], lactate+ketones+ethanol [LKA], and endogenous [END, no substrate added]) measured for half the individuals at 12°C and half at 28°C.

Heart and brain tissues were collected after measuring MR_cardiac_ and stored in chaotropic buffer for mRNA expression analysis. The mRNA data includes tissue specific expression counts for single mRNAs and tissue specific module mRNA expression (ME) from a whole genome co-expression network analysis (WGCNA, v1.70-3, (Langfelder and Horvath 2008; Drown et al. 2022)). The WGCNA approach summarizes mRNAs with correlated expression into co-expression modules, calculates a principal component of module expression for each individual (ME), and assigns a rank to single mRNAs within the module (module membership) based on their correlation to the ME. Here, we use the first principal component of module expression (ME) as a multivariate molecular level phenotype that may be predicted using genotype likelihoods. In addition, we examined association between genotype likelihoods and the top 10 mRNAs for each module (based on module membership).

### 1. Association studies

All results are reported from the score test conducted in ANGSD using -doAsso 2 with filters -minHigh 4, -minCount 1. These filters are relaxed from the default filters (default: - minHigh 10, -minCount 10) and allowed us to examine associations for all traits with reduced sample size requirements. The sample size for each association can be found in Table S2. For all phenotypes, acclimation temperature was included as a covariate. For SMR, CT_max_, and MR_cardiac_, acclimation order (individuals were acclimated to 12°C then 28°C or 28°C then 12°C) was included as an additional covariate. For SNP associations to mRNA expression, heart and brain mRNA expression were examined as separate phenotypes. P-values for genotype to phenotype associations were corrected for multiple testing using the Benjamini-Hochberg approach (Benjamini and Hochberg 1995) and significant associations identified as those with a corrected p-value <0.05. To examine patterns among independent SNPs, in cases where SNPs associated with the same phenotype were within 500 bp of each other, we pruned SNPs to keep the most significant SNP for each association and removed any within 500 bp of that SNP.

### 2. Annotation of significant SNPs

A bed file was generated from a SNP list using the genomic region for the SNP as the SNP location – 1 bp: SNP location. Bedtools intersect was used to obtain annotation information from the .GTF file for the current *F. heteroclitus* genome.

### 3. Statistical analysis

Data visualization and statistical analyses were conducted in R (v 4.0.5). An annotated script is available on Github (https://github.com/mxd1288/Genotypic_drivers.git). Association tests were carried out using ANGSD, as described above, and the likelihood ratio test values used to calculate p-values for each SNP to trait association calculating using a one-sided non-central chi-squared distribution (pchisq in R). P-values were corrected within each set of trait associations using p.adjust in R with the Benjamini and Hochberg method (Benjamini and Hochberg 1995).

## Contributions

Data collection, analysis, visualization, and initial draft manuscript writing MKD. Manuscript editing MFO and DLC. Funding from the National Science Foundation to MFO and DLC.

## Acknowledgements

The authors recognize Moritz Ehrlich and Amanda DeLiberto for assistance in Fall 2018 field collections and useful discussion of lcWGS analysis and genomic association studies. Amanda DeLiberto and Samantha Sierra-Martinez also purified the Tn5 enzyme used in library preparation.

## Tables and Figures

**Figure S1:**
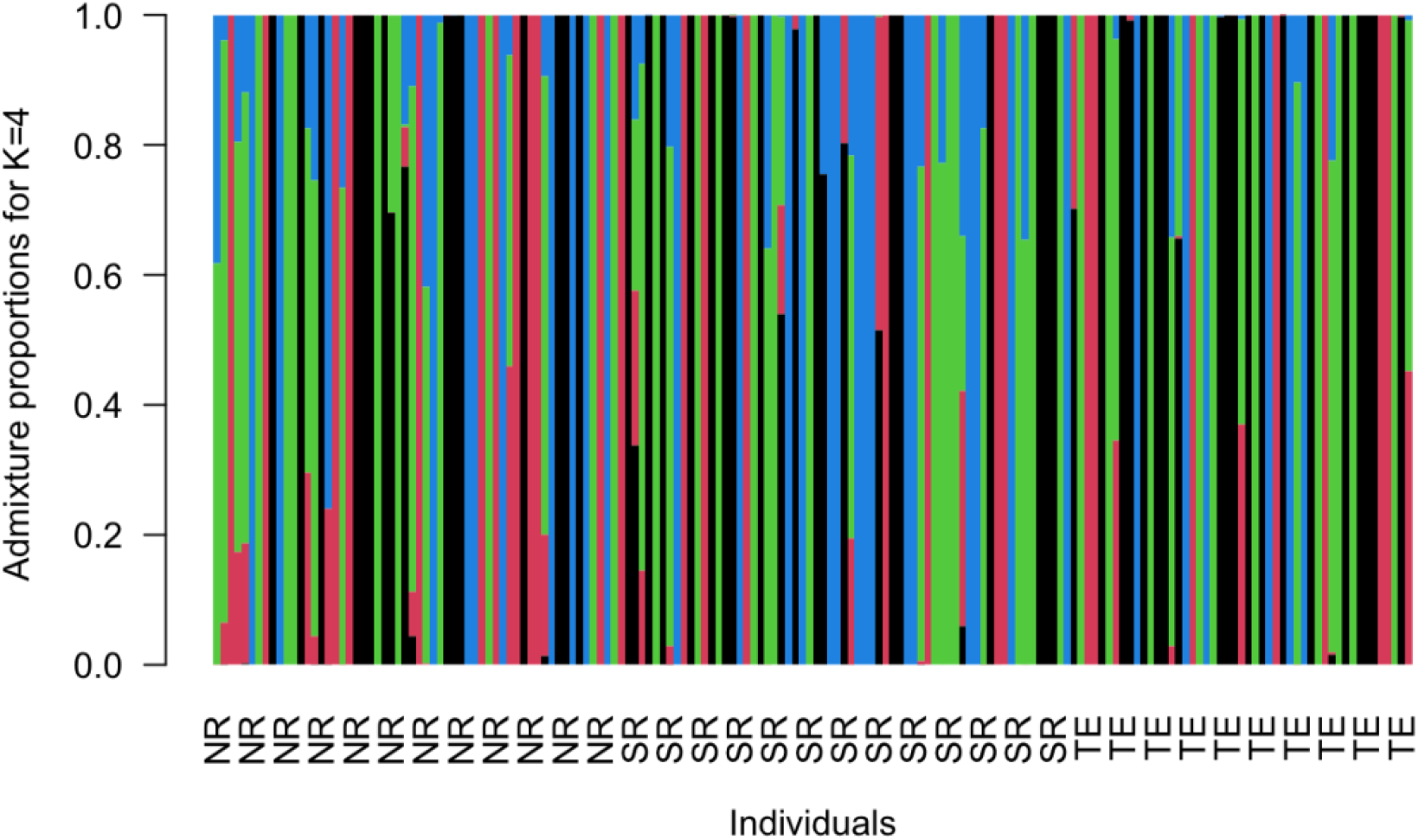
No evidence in admixture analysis of population structure among samples. The best K=4 based on log-likelihood probability for 172 individuals collected from three population in fall 2018: North Reference (NR), South Reference (SR), and Thermal Effluent (TE) populations.

**Figure S2:**
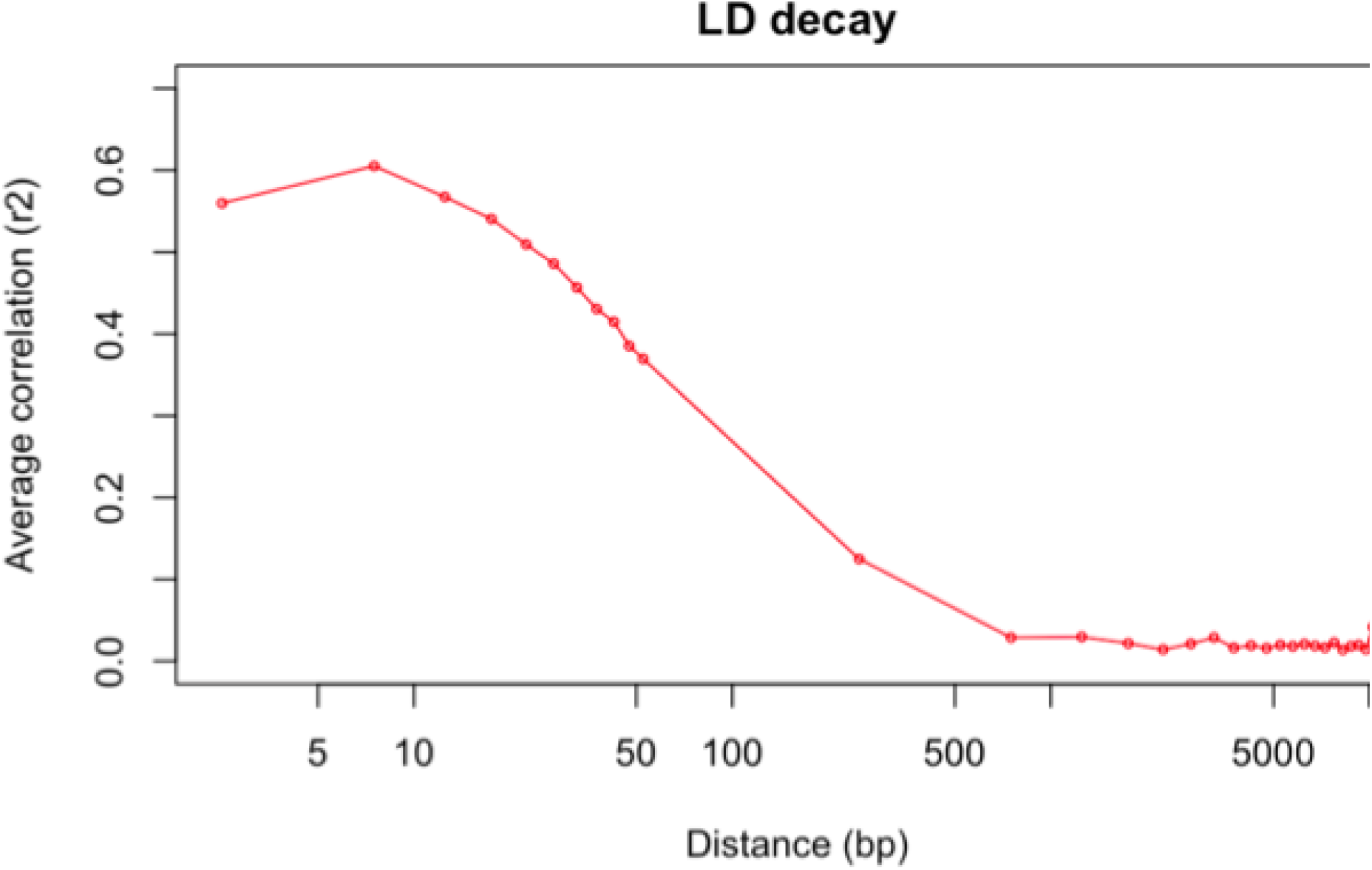
Decay of linkage disequilibrium. Linkage among SNPs decays to an average correlation axis) of <0.1 within 300bp.

**Figure S3:**
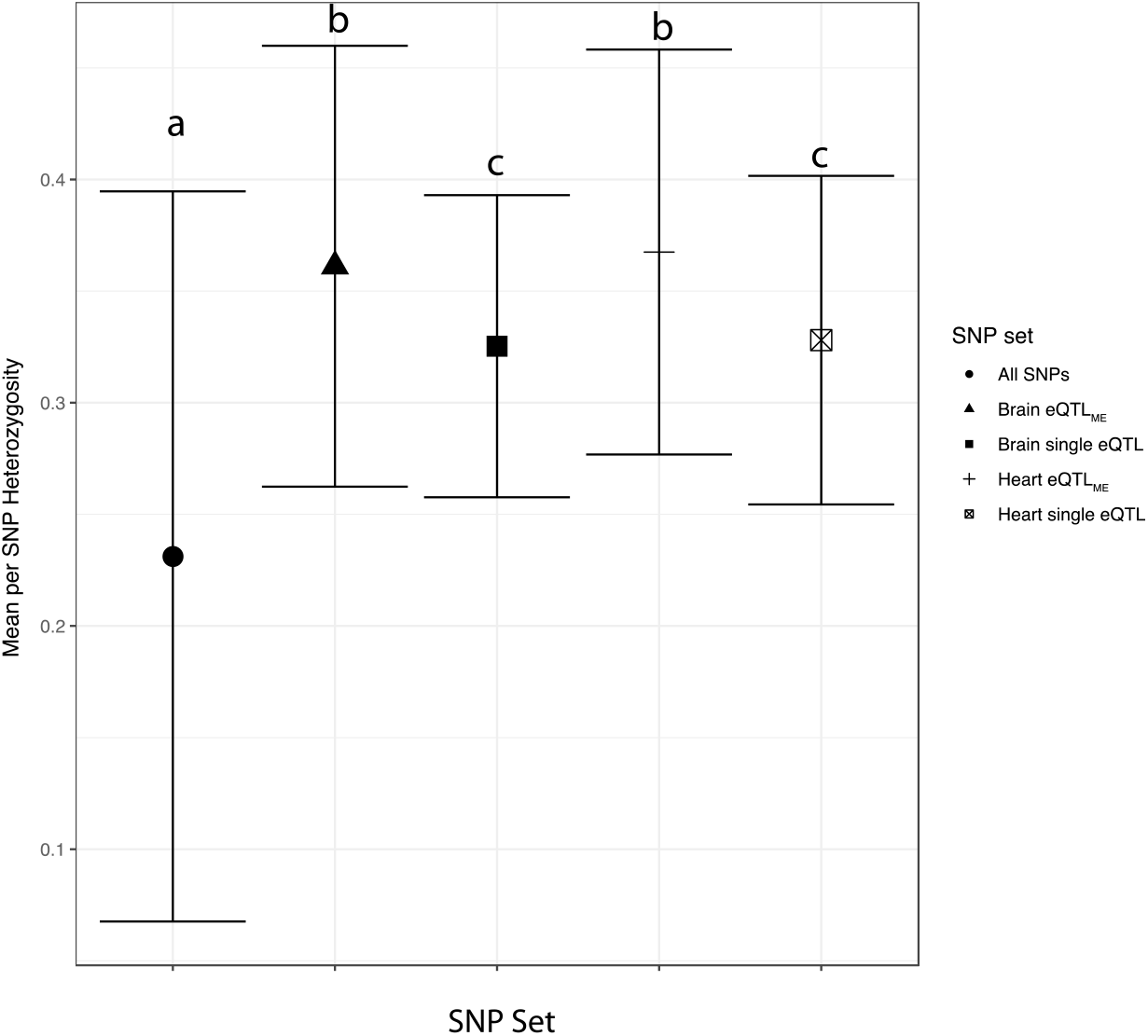
Heterozygosity among all single nucleotide polymorphisms and eQTL sets. Heterozygosity was significantly higher for SNPs identified as eQTL_ME_ and eQTLs when compared to all SNPs. Heterozygosity was significantly higher for eQTL_ME_ compared to eQTLs with no difference between tissues.

**Figure S4:**
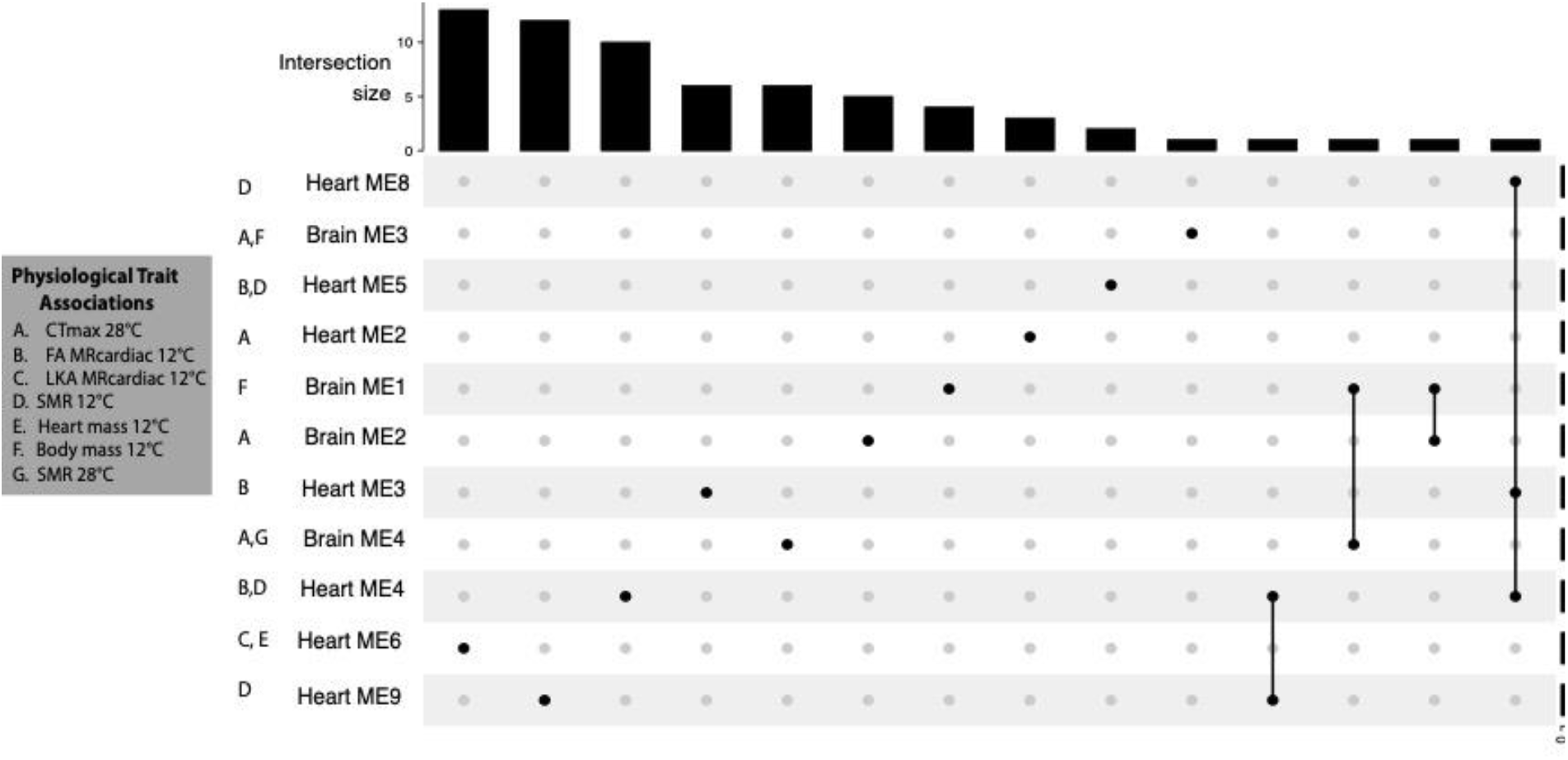
Overlap among eQTL_ME_ for modules correlated to physiological traits. Genetic control of modules that explain physiological trait variation appears independent with few shared SNPs in any eQTL_ME_ set. The exceptions are a single shared eQTL_ME_ between brain ME1 and brain ME4 (no shared trait correlation), between brain ME1 and brain ME2 (no shared trait correlation), and among heart ME3, heart ME4, and heart ME8 (shared correlation of ME3 and ME4 with 12°C FA MR_cardiac_ and shared correlation of ME4 and ME8 with 12°C SMR).

